# Spatial Perturb-Seq: Single-Cell Functional Genomics within Intact Tissue Architecture

**DOI:** 10.1101/2024.12.19.628843

**Authors:** Kimberle Shen, Wan Yi Seow, Choong Tat Keng, Daryl Lim Shern, Ke Guo, Amine Meliani, Irfan Muhammad Bin Hajis, Kok Hao Chen, Wei Leong Chew

## Abstract

We develop Spatial Perturb-Seq, an *in vivo* CRISPR technology that interrogates multiple genes within single cells of intact tissues. We apply Spatial Perturb-Seq to screen risk genes for neurodegenerative diseases in the mouse brain, uncovering cell autonomous and cell-cell microenvironmental effects within the spatially intact tissue. Spatial Perturb-Seq functionally screens multiple genes *in situ* and *in vivo*, identifying candidate genes underlying dysregulated neuronal intercellular communication pathways in Lrp1 signalling and ephrin-Eph receptor interactions.

## Main

Perturb-seq, which combines pooled CRISPR screens with single-cell RNA-sequencing (scRNA-seq), has dramatically increased the scale of functional genomics studies^1,2^. Several studies have showed feasibility of perturb-seq *in vivo*^3,4,5^, in a system where cells maintain their native gene expression patterns. However, spatial organization and cell-cell interactions are not preserved. Perturb-map^6^, relying on protein barcodes detected by imaging methods, has been used to identify genetic factors influencing the spatial organization of solid tumours. Similarly, optical pooled screens allow perturbations to be linked to image-based phenotypes but require extensive technical expertise and specialized microscopic setups^7,8,9,10^. These prior approaches uncouple the CRISPR perturbation detection platforms from the transcriptomic sequencing used for cell phenotype identification. In contrast, a spatial functional genomics technology that uses scalable nucleotide barcodes would allow the endogenous cell phenotype, exogeneous perturbations, and spatial localizations to be concurrently identified with a single sequencing technology. Here, we developed Spatial Perturb-Seq, an *in vivo* Perturb-seq technology that interrogates genetic functions directly in single cells in their native tissue. Spatial Perturb-Seq simultaneously measures whole transcriptomes at single-cell resolution (and therefore cell type), CRISPR barcodes (linked to gene perturbation), spatial coordinates, and cell-cell interactions (microenvironment) - all through a single Stereo-seq^11^ run.

We demonstrate Spatial Perturb-Seq by simultaneously interrogating 18 genes within the same animals, using a pooled barcoded CRISPR-gRNA AAV library (Extended Data Table 1) delivered intracranially into the hippocampus of Cas9-expressing mice^12^ at low multiplicity of infection (MOI). Intentional sparse editing ensures perturbed cells are surrounded by non-edited cells, allowing isolation of cell autonomous and non-cell autonomous (i.e. microenvironmental) effects. Most neighbours of perturbed cells are barcode-negative as designed (Extended Data Fig. 7). We used AAV over the more commonly used lentivirus, to avoid possible genotoxic integration^13^ and template switching events that uncouple barcodes from their respective gRNAs^14^. Tropism of the AAV-PHP.eB serotype used here was indeed highest in neurons^15^, with high tropism also observed for astrocytes and oligodendrocytes, but not for microglia and endothelial cells (Extended Data Fig. 1a-e). This potentially reflects the similar lineages of neurons, astrocytes and oligodendrocytes being derived from neural stem cells.

We first profiled the injected mouse brains with conventional single-cell *in vivo* Perturb-seq using 10X scRNA-seq (Extended Data Fig. 1a-c). Among the different cell types, we recovered predominantly oligodendrocytes, with 11% of oligodendrocytes being positive for barcodes and an average of 26 oligodendrocytes per perturbation (Extended Data Fig 1d-g). Neurons are underrepresented, as they are fragile during the cell dissociation process used^16^. Each AAV delivers 3 gRNAs targeting the same gene, because gene editing efficiency is increased using multiple guides compared to single guides (Extended Data Fig. 2a), and to reduce perturbation failure due to non-functional guides. In silico off-target prediction for all 54 guides identified only 6 sites in the mouse genome with <3 mismatches. All sites with <4 mismatches are shown in Extended Data Table 2. To confirm CRISPR-Cas9 activity in Cas9 mice, we transfected gRNAs into BMDMs derived from Cas9 mice and achieved high editing rates (Extended Data Fig. 2b). We also verified *in vivo* editing in the tissue of injected mice, observing an average of 0.15% indels for each locus genotyped in the single cell lysate (Extended Data Fig. 2c, d), close to the observed mean of 0.13% of sequenced single cells containing each barcode. Mutational profiling showed distinct indels between 1-10 bp (Extended Data Fig. 2e), consistent with precise on-target CRISPR/Cas9-mediated editing at the target sites. This suggests that cells delivered with barcoded CRISPR gRNAs are largely edited at the genomic loci sequenced, consistent with editing efficiencies of single gRNAs in Cas9-mice^3^. The consistency between the intentional low fraction of cells harbouring individual barcodes and the fraction of cells exhibiting targeted genetic perturbations, together indicate that barcode detection by sequencing is a reasonable proxy for inferring linked genetic perturbations.

Focussing on oligodendrocytes, we recovered known oligodendrocyte subtypes: MOL1, MOL2, MOL5/6, and DA_Ifn^17^ (Extended Data Fig. 3a, b). The proportion of barcode-positive MOL1 oligodendrocytes was significantly higher than barcode-positive MOL5/6 oligodendrocytes (Extended Data Fig. 3c, d), suggesting that injection injury and AAV transduction could steer oligodendrocytes to a less mature, MOL1 state, consistent with reports showing MOL1 enrichment at injury sites^18^. It is also possible that AAV-PHP.eB has stronger tropism for MOL1 oligodendrocytes. Oligodendrocytes are responsible for making and maintaining the myelin sheath, a process that involves lipid metabolism, which we examined through the expression of two sets of genes in oligodendrocytes: myelin genes (*Mobp, Mog, Opalin, Plp1, Mbp, Cnp, Mag, Mal*) and lipid metabolism genes (*Srebf1, Hmgcr, Scd1, Scd2, Acaca, Dgat1, Cpt1a, Elovl6*). *Olig2-KO* does not significantly reduce the expression of canonical mature myelin genes (Extended Data Fig. 3e), consistent with its role in oligodendrocyte specification but not maintenance^19^. *Rraga* and *Flcn* perturbations cause slight downregulation of myelination genes in mice (Extended Data Fig. 3e), suggesting that the Rraga-Flcn-Tfeb pathway could be a promising target for promoting remyelination. This is consistent with Flcn and Rraga being required for myelination in zebrafish^20^, even though their function in mammalian myelination has not been recognized. In addition, Fasn, a key regulator of fatty acid metabolism, is required for the expression of lipid metabolism genes in oligodendrocytes (Extended Data Fig. 3f).

Spatial Perturb-Seq provides spatial data that reveals biological interactions beyond conventional single-cell Perturb-Seq. Applying *in vivo* Spatial Perturb-Seq with Stereo-seq (Fig. 1a), we obtained spatial sequencing data for 229,775 cells over 3 experiments and 4 mice (Extended Data Fig. 5a and Extended Data Table 3). A subsection of stereo-seq data highlights individual cells with distinct colours for each barcode (Extended Data Fig. 4a). All 18 barcodes are represented in the sequencing data (Extended Data Fig. 4b, c). The libraries were prepared without barcode-specific PCR-enrichment to avoid quantification bias. Nonetheless, intentional barcode enrichment did not drastically increase the number of barcode-positive cells, but increased the detected levels of barcodes, indicating that the sequencing saturation is sufficient (Extended Data Fig. 5b). The dataset is then processed with DeepCell^21^ for cell segmentation (Extended Data Fig. 5c), where an average of 462 and 773 expressed genes were detected per cell and per neuron, respectively (Extended Data Table 3). We recovered neurons, RBCs, oligodendrocytes, microglia, astrocytes, choroid plexus and endothelial cells (Fig. 1b-f) with spatial patterns consistent with gene expression modules (Extended Data Fig. 5d, e). The consistency in our manual annotation and spatial hotspot analyses of gene modules, provides confidence in the data quality and cell segmentation.

**Fig. 1:**
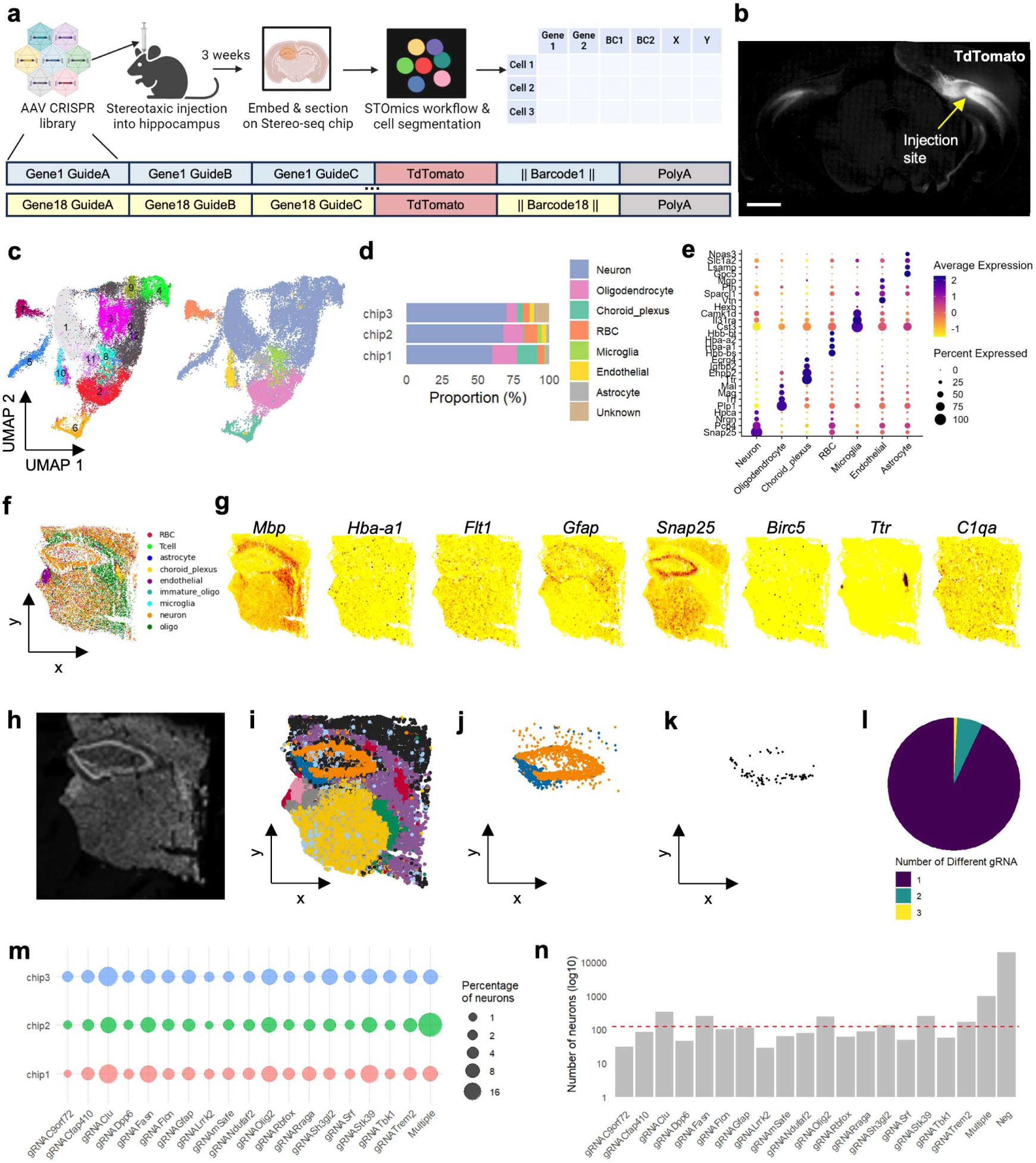
*In vivo* spatial perturb-seq of the adult mouse brain by intracranial injection of a barcoded AAV library. (a) Schematic of the *in vivo* spatial Perturb-seq pipeline, and depiction of components in our AAV-PHP.eB library, showing 3 gRNAs, TdTomato and 1 barcode in each member. (b) Confocal imaging of a coronal section of a mouse brain following stereotaxic injection into the hippocampus, showing that transduction spread is limited to the hippocampus. Scale bar: 1000 µm. (c) UMAP projection of cells from chip 1 following Stereo-seq, showing recovery of neurons, oligodendrocytes, choroid plexus cells, RBCs, microglia, endothelial cells, and astrocytes. Left: coloured by cluster number. Right: coloured by cell type. (d) Similar proportions of cell types were recovered for each chip, with the majority cell type being neurons as expected. (e) Dot Plot showing expression of cell-type enriched genes for each cell type. (f) Spatial scatter of annotated mouse brain cells. (g) Spatial gene plot of canonical cell-type markers major cell populations (Microglia: C1qa; Oligodendrocytes: Mbp; Endothelial cells: Flt1; Neurons: Snap25; Astrocytes: Gfap, Choroid Plexus: Ttr; RBC: Hba-a1; Immature oligodendrocytes: Birc5). (h) ssDNA image for cellular localization on the Stereo-seq chip. (i) Spatial scatter following BANKSY clustering with lamba 0.2, showing clusters recovered, which show strong alignment between the two sections. (j) Selection of CA and DG neurons in the hippocampus by focusing on BANKSY clusters 3 and 10. (k) Subset of barcode-positive cells from (j). (l) Piechart representation of the number of unique gRNA barcodes among all gRNA barcode positive cells, showing a small proportion of barcode-positive cells with multiple barcodes detected. (m) Dot Plot showing percentage of barcode-positive neurons among hippocampal neurons in (j) for each chip. (n) Total number of cells for each gRNA across all 3 chips. Min: 29, max: 330, mean: 122. (f) – (l): Data from chip 1.

Spatial Perturb-Seq enables the interrogation of cell-autonomous versus microenvironment effects of each gene perturbation within native intact tissues. We employed BANKSY^22^ to select the region of interest, through adjusting the lambda parameter to vary the cells’ embedding. At low lambda values, BANKSY functions in single cell-typing mode, whereas at high lambda values, it identifies spatial domains (Extended Data Fig. 6). We used lambda 0.2 to achieve the goal of spatially informed cell-typing to identify hippocampal neurons (Fig. 1g-i). Among hippocampal neurons, all 18 barcodes were detected, with most barcode-containing cells containing only 1 unique barcode as designed (Fig. 1j-m). Cells with multiple barcodes (12.5% on average) were excluded from the analysis. Among all cells profiled by Stereo-seq, 2.1% are barcode-positive (Extended Data Table 3), closely aligning with the 2.4% observed in the 10X dataset, suggesting comparable barcode detection sensitivity.

To compare the effects of the different perturbations, we quantified the number of DEGs (p < 0.05, lfc > 0.5) for each perturbation compared to control, and their average effect size (Fig. 2b). As expected, perturbations cause a stronger cell autonomous effect, compared to their influence on the microenvironment (defined as 15 closest neighbouring cells). A caveat of this analysis is the assumption that the niche is uniform, whereas there may be spatial and microdomain-specific differences. Cfap410 perturbation caused the highest number of DEGs in its non-perturbed cellular neighbourhood, consistent with its role as a player in cilia function and synaptic plasticity in neurons. Overall, there are fewest DEGs in Rraga-KO neurons, potentially due to redundancy between *Rraga* and *Rragb* in neurons^23^. The top 5 significantly upregulated genes for each perturbation were distinct among the 18 perturbations (Fig. 2c), highlighting the specificity of transcriptomic changes induced by each genetic KO.

**Fig. 2:**
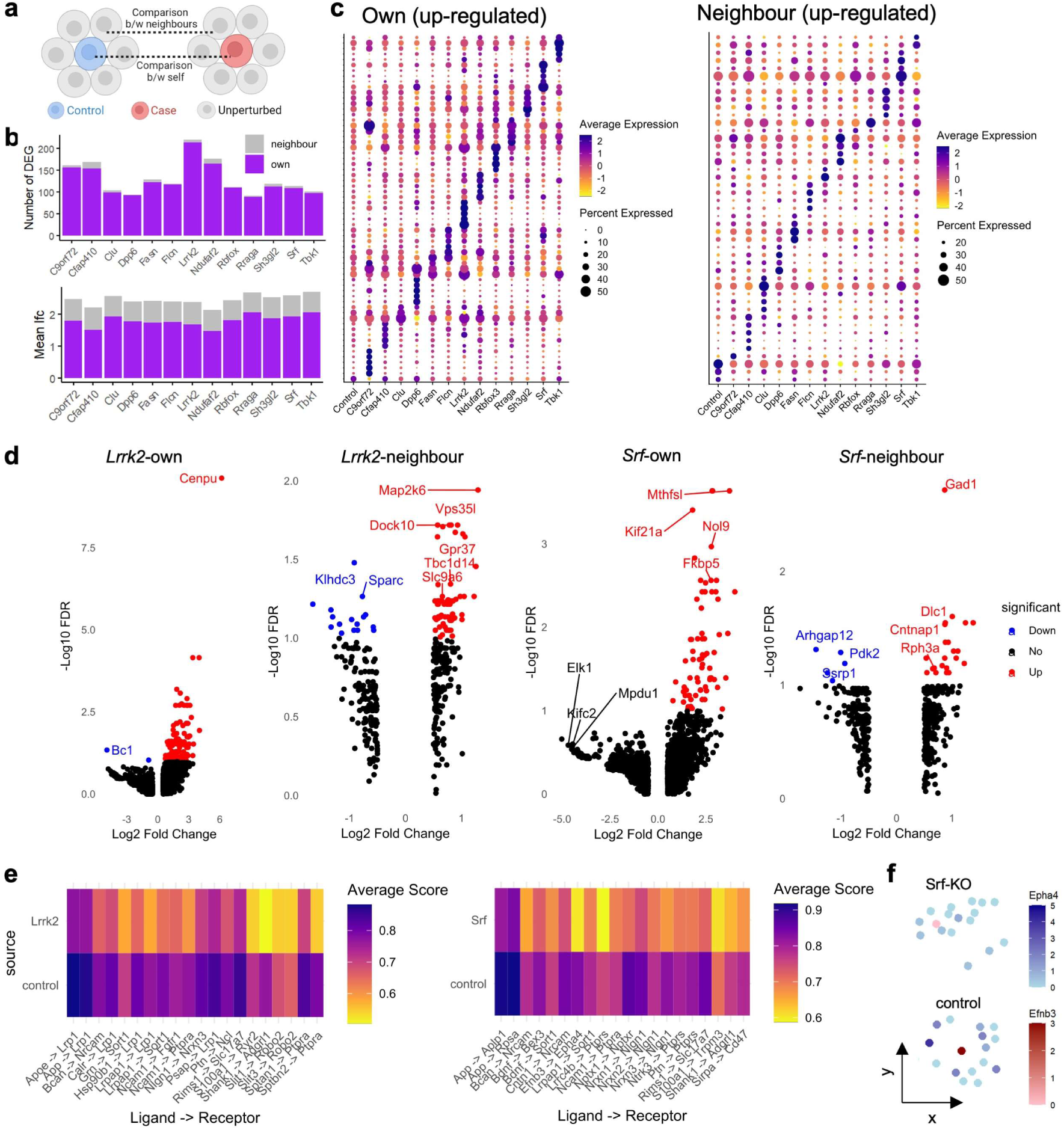
Analysis of cell-autonomous and non-cell autonomous effects of perturbations and cell-cell communication in hippocampal neurons. (a) Schematic showing two different comparisons that can be made – cell autonomous and non-cell autonomous, i.e. 1) transcriptomes of perturbed cells (own). 2) transcriptomes of the wildtype neighbours of perturbed cells (neighbours). (b) Top: Number of DE genes (lfc > 0.5, pval < 0.05) for each perturbation (own, neighbours) compared to the control group. Bottom: Average lfc for the DE genes. (c) Dot Plot showing top DE genes for each of perturbation, compared to all other perturbations. Left: Own transcriptome. Right: Neighbour cells transcriptome. (d) Volcano plots showing DE genes (lfc > 0.5, fdr < 0.05) for Lrrk-KO and Srf-KO own and neighbour cells. Only lfc > 0.5 genes are shown. (e) Heatmap visualizing ligand-receptor expression for top 20 differently-expressed ligand-receptor pairs, comparing communication scores between Lrrk2-KO or Srf-KO neurons and their wildtype neighbours vs communication between control neurons and their wildtype neighbours. (f) Spatial plot of each source (ligand-expressing) cell in red and their 15 wildtype neighbour (receptor-expressing) cell in blue, showing expression of *Epha4* ligand in perturbed cells and *Efnb3* receptor in neighbour cells. Top: Srf-KO. Bottom: control.

Knock out of *Lrrk2* (n = 29 cells) led to 213 DEGs observed (p < 0.05, lfc > 0.5), the highest number amongst all perturbed genes (Fig. 2b-d). *Lrrk2* is a key player in neuronal signalling and Parkinson’s disease pathology^24,25^. *Bc1*, a lncRNA found in dendrites that regulates translation of specific mRNAs in synapses^26^, is downregulated upon *Lrrk2* KO. Among the microenvironmental changes induced by *Lrrk2* KO: downregulation of *Sparc*; upregulation of *Vps35l* (Vps35 is known to functionally interact with Lrrk2^27^), *Dock10* (involved in dendritic spine formation^28^), and *Gpr37* (known to be associated with PD^29^). The microenvironmental effects stemming from *Lrrk2* KO can potentially underlie Lrrk2-mediated pathology in neuronal signalling and Parkinson’s disease.

Another striking finding is that from *Srf* (n = 50 cells), a transcription factor regulating genes involved in neuronal growth and synaptic plasticity^30^. Among the genes dysregulated upon Srf-KO, most are target genes identified through ChIP-seq data from the ENCODE Transcription Factors Target dataset^31^, though some did not reach the fdr threshold of < 0.05. In the Srf-KO microenvironment, *Arhgap12* (involved in cytoskeletal and actin dynamics^32^) and *Ssrp1* (co-activator of Srf^33^) were downregulated, *Gad1* (involved in the synthesis of neurotransmitter GABA^34^) was upregulated.

We dived deeper into cell-cell communication, as it is essential in the function of neurons. To ensure accuracy of cell-cell communication analysis, we studied cells by keeping to the same neuronal and spatial niches, as ligand-receptor communication occur among neighbouring cells in proximity. We used the LIANA receptor-ligand framework^35^ to interrogate the dataset and prioritize biologically important ligand-receptor pairs (Fig. 2e). We observed a 19% reduction in the SCA score for Lrp1 signalling in the cellular neighbours of Lrrk2-KO neurons, compared to control neurons. Lrp1 mediates important neuronal functions including α-synuclein uptake^36^, suggesting a possible mechanistic link between Lrrk2 activity and PD pathogenesis. The data showed a reduction in Nlgn1 signalling in Srf-KO neighbours, highlighting the importance of Srf in synaptic function. To visualize these cell communication networks, individual cells showing Efnb3-Epha4 signalling in Srf-KO and control-KO microenvironments is shown on a spatial plot (Fig. 2f). Source cells and their 15 closest neighbours (target cells), are shown according to their XY coordinates, with expression levels of the ligand in red (source cells) and expression levels of the receptor in blue (target cells). These analyses highlight the ability of Spatial Perturb-Seq to interrogate genetic determinants of intercellular cell communication signalling pathways within the native tissue.

We present Spatial Perturb-Seq, which enables the interrogation of genetic perturbations on cells and their microenvironment directly in the tissue architecture with single-cell resolution and whole transcriptome coverage. This new ability to segment a genetic perturbation’s autonomous and non-cell autonomous effects reveals insights into cell-cell communication that would otherwise be lost via conventional dissociated single cell sequencing. To assess compatibility of our system with an alternative nucleotide-based detection system, we performed RNA FISH against our nucleotide barcodes and found high specificity FISH spots (R2 = 0.78) (Extended Data Fig. 9), demonstrating compatibility with probe-based spatial transcriptomics platforms such as MERFISH^37^ or CosMx^38^. Further introduction of cell-type specific barcode expression or sequential injections of different barcodes could endow capabilities for spatial lineage tracing^39^ and spatial molecular timing^40,41^. The primary bottlenecks of Spatial Perturb-Seq are cost of sequencing as both perturbed cells and their unperturbed neighbours must be sequenced, and the naturally limited cell numbers in a tissue niche. However, the steady decline in sequencing costs will enable larger sample sizes to be sequenced to increase statistical power. Power analysis done with powsimR^42^ to assess the statistical robustness of our platform showed FDR <10% and TPR >60-80% for 125-625 cells per perturbation (Extended Data Fig. 8), demonstrating suitability of multiplexed experiments using this platform. As is the case with most spatial technologies, the accuracy of cell segmentation underpins the accuracy of Spatial Perturb-Seq, though significant advancements are being made in the field. Spatial Perturb-Seq unlocks a strategy to functionally interrogate genes at single-cell resolution within intact tissues, enabling the profiling of perturbation effects in both the target cells and their wildtype microenvironment.

## Methods

### Experimental methods

#### Mice

All animal work was performed under the guidelines of the Institutional Animal Care and Use Committee (IACUC protocol #211646). Mice were housed on a standard light cycle under pathogen-free conditions. Female Rosa26-CAG-Cas9 mice (JAX 024858) aged 8-16 weeks were used.

#### Stereotaxic injections

Animals are anesthetized with ketamine/xylazine (injected intraperitoneally) and placed on a thermostatically controlled heating pad. Animals were checked for sedation. Hair was removed from the incision site on the scalp with a shaver and the area was cleaned with an alcohol wipe. Corneas were protected with an ophthalmic lubricating ointment. Animals were then positioned in a stereotaxic head holder with the skull firmly barred. An incision on the scalp was made and the scalp reflected over the brain region of interest. For all experiments, the left hippocampus was targeted with the following stereotaxic coordinates relative to the bregma: AP −1.7, ML +1.6, DV −1.8. A handheld dental drill was used to drill a <1mm diameter hole in calcified bone in the AP, ML region of interest. The cannula was inserted into the region of interest and the 5 x 10^8 viral particles of AAVs were injected at a rate of 2-5nl/second in a maximal volume of 1.5ul, so as not to cause tissue injury. Following injection, the cannula is withdrawn after 5 minutes. The incision on the scalp was closed with Vetbond Tissue Adhesive. Atipamezole was administered intraperitoneally for anaesthesia reversal and Buprenorphine was administered subcutaneously for pain relief. Following which, animals were placed on a thermostatically controlled heating pad for recovery and monitoring. For 3 days post-surgery, Buprenorphine is administered for pain relief, and animals were monitored for signs of distress and wound inflammation. Mice were kept for 3 weeks under standard conditions before being sacrificed by CO2 asphyxiation for tissue extraction and processing.

#### gRNA library design

We designed three guides each for 18 loci - 17 genes and a safe harbour locus (*mSafe*) as a control (54 guides in total). The three guides for the same locus are linked to the same barcode on one vector, to facilitate detection of the genetic knockout via a single unique barcode. Each vector contains TdTomato, with 18 vectors in total. We focused our library to target genes associated by GWAS. From the NHGRI-EBI Catalog of human genome-wide association studies^43^, we chose GWAS hits associated with Alzheimer’s Disease (https://www.ebi.ac.uk/gwas/efotraits/MONDO_0004975) (CLU, LRRK2 NDUFAF2, RBFOX1, TREM2), Amyotrophic Lateral Sclerosis (https://www.ebi.ac.uk/gwas/efotraits/MONDO_0004976) (C9orf72, CFAP410, DPP6, TBK1), and Parkinson’s disease (SH3GL2, STK39) (https://www.ebi.ac.uk/gwas/efotraits/MONDO_0005180). These genes have mouse orthologs and are expressed in the mouse brain. We also included 6 positive control genes (Fasn, Flcn, Gfap, Olig2, Rraga, Srf) and a safe harbour site. With these 18 genes/loci of interest, we designed three gRNAs each using the online tool CHOPCHOP (https://chopchop.cbu.uib.no/). Best-scoring guides within the first few exons were selected.

#### Cloning of gRNAs plasmid pool

The plasmid pAAV-CAG-tdTomato (codon diversified) was ordered from Addgene (#59462). The plasmid backbone (2ug) was digested with HindIII/SalI (NEB) for 1h at 37°C followed by an inactivation step for 20 mins at 80°C. DNA fragments consisting of both the overlapping sequences with the digested pAAV-CAG-tdTomato backbone and the 450 bp barcodes were ordered from IDT. PCR reactions were used to amplify the DNA fragments. The Gibson assembly reaction was set as follows: 100 ng of digested plasmid backbone, 10ng of amplified DNA fragment, 10 µl NEBuilder HiFi DNA Assembly Master Mix (NEB, E2621L) and H2O up to 20 µl total reaction. The reaction was incubated for 1 h at 50°C. 5ul of the Gibson reaction was used for transformation using Stbl3 cells (Thermo Fisher, Cat#C737303). Sequencing reactions (Biobasic) were used to confirm the correct insertion of the barcodes. Following barcodes cloning, three gRNAs were designed for each locus-of-interest to be inserted into each barcoded plasmid. Golden gate cloning strategy was employed to arrange the three gRNAs in tandem, each driven by a U6 promoter. The gRNAs were then inserted into the NdeI-digested pAAV-CAG-tdTomato using Gibson assembly.

#### AAV production and purification

AAVs were generated in-house as per previously reported^44^. Briefly, AAVs were packaged via a triple transfection of 293AAV cell line (AAV-100, Cell Biolabs, San Diego, CA, USA). The cells were plated in a HYPERFlask ‘M’ (Corning) in growth media which consists of DMEM + glutaMax + pyruvate + 10% FBS (Thermo Fisher), that was supplemented with 1X MEM nonessential amino acids (Gibco). Transfection was done when the cells were between 70% and 90% confluent. Media was replaced with fresh pre-warmed growth media before transfection. For each HYPERFlask ‘M’, 200 μg of pHelper (Cell Biolabs), 100 μg of pRepCap encoding capsid proteins for serotype DJ or 2], and 100 μg of pZac-CASI-GFP or pZac-CMV-CasRx gRNAs were mixed in 5 ml of DMEM, and 2 mg of PEI “MAX” (Polysciences) (40 kDa, 1 mg/ml in H2O, pH 7.1) was added for PEI: DNA mass ratio of 5:1. The mixture was incubated for 15 min before being transferred to the cell media drop-wise. The day after transfection, the media was changed to DMEM + glutamax + pyruvate + 2% FBS. 48–72 h after transfection, the cells were harvested by scrapping or dissociation with 1X PBS (pH7.2) + 5 mM EDTA and then pelleted at 1500 g for 12 min. Cell pellets were resuspended in 1–5 ml of lysis buffer (Tris HCl pH 7.5 + 2 mM MgCl + 150 mM NaCl), before freeze-thawing thrice using a dry-ice-ethanol bath and a 37°C water bath. Cell debris was clarified via centrifuging at 4000 g for 5 min, and the supernatant was collected. To remove the unpackaged nucleic acids, the collected supernatant was treated with 50 U/ml of Benzonase (Sigma–Aldrich) and 1 U/ml of RNase cocktail (Invitrogen) for 30 min at 37°C. Next, the lysate was loaded on top of a discontinuous density gradient that consisted of 6 ml each of 15%, 25%, 40%, and 60% Optiprep (Sigma– Aldrich) in a 29.9 ml Optiseal polypropylene tube (Beckman–Coulter). The tubes were then ultra-centrifuged at 54,000 rpm, for 1.5 h at 18°C, using a Type 70 Ti rotor. The 40% fraction was extracted and dialyzed with 1X PBS (pH 7.2) supplemented with 35 mM NaCl, using Amicon Ultra-15 (100 kDa MWCO) (Millipore). qPCR was then carried out with the ITR-sequence-specific primers and probes, using the ATCC reference standard material 8 (ATCC) as a standard, to determine the titres of the purified AAV vector stocks.

#### Tissue dissection, dissociation and FACS

After mice were sacrificed by CO2 asphyxiation, intracardial perfusion with ice-cold PBS was carried out to remove blood cells. Brains were dissected and either embedded in OCT for sectioning or dissociated into single cells. Dissociation of brain tissue was done with the Neural Dissociation Kit (Miltenyi Biotec) following manufacturer’s instructions. Briefly, brain tissue of interest was minced with a razor blade followed by enzymatic digestion at 37℃ and triturated. The cell suspension was then strained through a 70um cell strainer and centrifuged at 300g for 10min. The cell pellets were then re-suspended in Hibernate A with 10uM Calcien-AM (Invitrogen) and 1ug/ml Propidium Iodide (PI) (Stemcell), for FACS sorting. Live, single cells were selected by positive selection of Calcien-AM and negative selection of PI. Immediately after FACS sorting, cells were used for the generation of 10X 3’ chromium libraries and sequenced on Novaseq (Illumina).

#### Coverslip functionalization and sample preparation for FISH

Coverslip functionalization was carried out before use for tissue sectioning. Coverslips (Warner Instruments, cat. no. 64–1500) were cleaned in 1M KOH for 1 h before being rinsed thrice in Mili Q water. The coverslips were rinsed with 100% methanol before being functionalised in an amino-silane solution (3% vol/vol (3-aminopropyl) triethoxysilane (Merck cat no. 440140), 5% vol/vol acetic acid (Sigma, cat. no. 537020) for 2 min at room temperature. Following which, the coverslips were rinsed thrice with Mili Q water before being dried overnight at 47°C in an oven. The mouse brain sample was frozen in optimal cutting temperature compound and sectioned into 7 μm sections onto functionalized coverslips using a cryostat. The sections were fixed using 4% vol/vol paraformaldehyde in 1× PBS for 15 minutes, then rinsed with 1× PBS and stored at -80°C.

#### FISH

The encoding probes were diluted in a 10% hybridization solution that was composed of 10% deionized formamide (Ambion™ Cat: AM9342) (vol/vol), 1 mg ml-1 yeast tRNA (Life Technologies, cat. no. 15401-011) and 10% dextran sulfate (Sigma, cat. no. D8906) (wt/vol) in 2× SSC. A final concentration of 10-50 nmol per probe was used. Mouse brain sections were permeabilized with 70% ethanol overnight at 4°C. After permeabilization, samples were rinsed twice with 2× SSC. Next, encoding probe staining was performed and samples were stained for 16 h at 37°C. After encoding probe staining, samples were washed in 10% formamide wash buffer (10% deionized formamide in 2× SSC) at 37°C for 15 min, twice. The samples were rinsed thrice with 2× SSC, before DAPI (Sigma, cat. no. D9564) staining was carried out.

#### Microscopy for FISH

Microscopy and acquisition of fluorescent images was carried out on a custom-built microscope, constructed using a Nikon Ti2-E body. A Marzhauser SCANplus IM 130 mm × 85 mm motorized X–Y stage, a pco.edge 4.2 BI-USB Back Illuminated sCMOS camera, a custom fiber-coupled laser box from CNI laser for illumination, and a Nikon CFI Plan Apo Lambda 60× 1.4-n.a. oil-immersion objective (MRD01605) was used. The Nikon Perfect Focus system was used to maintain focus while imaging, and in each imaging cycle, one Z position was imaged for each field of view. Custom in-house software was used for acquiring the images. Imaging was carried out on a home-build microscope^45^ constructed around a Nikon Ti2-E body. We used a Marzhauser SCANplus IM 130 mm × 85 mm motorized X–Y stage, and an Andor Sona 4.2B-11 sCMOS camera, and imaging was done with a Nikon CFI Plan Apo Lambda 60× 1.4-n.a. oil-immersion objective. Lasers used for illumination were as follows: 2RU-VFL-P-500-592-B1R (500 mW), 2RU-VFL-P-1000-647-B1R (1000 mW), 2RU-VFL-P-500-750-B1R (500 mW) (MPB Communications), and Coherent Obis 405 100-mW laser. We use an exposure time of 1 s for imaging, and focus was kept using the Nikon Perfect Focus System (PFS).

Samples were mounted onto the microscope stage via use of a flow chamber (Bioptechs, cat. no. FCS2). This allowed buffer exchange to be carried out via use of a computer-controlled fluidics system (Cite2). The sample was stained at room temperature for 15 min with readout probe solution via buffer exchange, prior to imaging. The readout probe solution consists of 10 nM of each fluorescently labelled readout probe, 10% deionized formamide (vol/vol) and 10% dextran sulfate (wt/vol) in 4× SSC. After hybridization, 2× SSC was flowed in before a rinse with 10% formamide wash buffer. 2× SSC flowed again before imaging buffer. The imaging buffer contains 2× SSC, 10% glucose, 50 mM Tris-HCl pH 8, 2 mM Trolox (Sigma, cat. no. 238813), 40 μg/ml catalase (Sigma, cat. no. C30), and 0.5 mg/ml glucose oxidase (Sigma, cat. no. G2133). Signal removal was done by washing the samples with a 55% formamide wash buffer containing 0.1% TritonX-100.

#### 10X library preparation and sequencing

Libraries from dissociated single cells were generated using Chromium Next GEM Single Cell 3’ Reagent Kits v3.2 (Dual Index) (10X Genomics). Cell numbers were counted by FACS and 20,000 cells per mouse was used to generate each library. Libraries were generated according to the manufacturer’s instructions, with 12 cycles of cDNA amplifications, 25ng of cDNA input and 14 cycles of sample index PCR. Barcode sequences were not amplified to avoid introduction of biasness and overamplification. Libraries generated had a fragment length of 400-450bp and were sequenced on an Illumina Novaseq (PE150).

#### iSeq library preparation and sequencing

In order to confirm the presence of indels in brain tissue after AAV transduction, 50k TdTomato+ cells from 4 brains were lysed with QuickExtract (Lucigen) according to manufacturer’s instructions. Uninjected brain tissue was used as WT control. Two rounds of PCR were used. For PCR1, genotyping primers were designed using Primer3 (https://primer3.ut.ee/) to amplify a 150-300bp region around the Cas9 cut site with 34 PCR cycles. The following sequences were added to the forward and reverse genotyping primers respectively, for the second round of amplification: CTTTCCCTACACGACGCTCTTCCGATCTNNNNNN, GGAGTTCAGACGTGTGCTCTTCCGATCT. NNNNN is used to introduce diversity at the start of the iSeq read to improve read quality. For PCR2 used to index samples, 6 PCR cycles were used.

For both PCR1 and PCR2, Q5 High Fidelity Master Mix (NEB) was used according to manufacturer’s recommended PCR temperatures and parameters. Indexed PCR products from PCR2 was pooled, ran on a gel, and gel purified using Wizard SV Gel and PCR Clean-Up System (Promega). DNA concentration was measured with Qubit HS dsDNA kit (Vazyme). The library was spiked with 2% PhiX library (Illumina) and sequenced on an iSeq 100 system.

#### Stereo-seq chip preparation

OCT blocks were stored at -80°C and equilibrated at -20°C for 2 h prior to sectioning. OCT blocks were cryosectioned at a thickness of 10 µm using a CM1950 cryostat. Following satisfactory QC results (RIN = 8.5-8.6), mouse brain coronal sections were collected on the Stereo-seq chip surface. Tissue sections were adhered to a Stereo-seq chip surface and was incubated at 37°C for 5 min. The tissues were then fixed in methanol and incubated at –20°C for 30 min. The tissue sections were then permeabilized at 37°C for 10 min and then washed with 0.1× SSC buffer. The RNA released from permeabilized tissues was captured using DNB probes and reverse-transcribed at 42°C for 3 hrs. After in situ reverse transcription, tissues were removed by tissue removal buffer. The chips were then incubated with 400 μL cDNA release buffer overnight at 55°C, and then washed once with 400 μL of 0.1× SSC buffer. The released cDNA was then collected and purified using 0.8× Ampure XP Beads. The cDNA was then amplified using the following PCR conditions 95°C for 5 min, 15 cycles of 98°C for 20 s, 58°C for 20 s, then 72°C for 3 min, and a final incubation at 72 °C for 5 min. PCR products were purified using 0.6× Ampure XP Beads and the concentrations of cDNA were quantified using a Qubit™ dsDNA Assay Kit.

#### Stereo-seq library preparation and sequencing

STOMICS libraries were constructed with the STOMICS Gene Expression kit. The cDNA was first checked for presence of barcodes by PCR. Briefly, 20ng of cDNA were fragmented at 55°C for 10 min. The fragmented products were then amplified using the following PCR conditions one cycle at 95°C for 5 min, 13 cycles at 98°C for 20 s, 58°C for 20 s and 72°C for 30 s, and lastly one cycle at 72°C for 5 min. The PCR products were purified using Ampure XP Beads for DNB generation and were finally sequenced (100 bp PE) using T7 sequencer.

### Computational methods

#### Barcode design

One of our goals is to design nucleotide barcodes long enough to be probed against, that are tolerated in cells. We wrote a script that can generated a set of barcodes with these features: having an appropriate length - long enough for probing against, but short enough to avoid inducing deleterious effects in the cell (450nt); not containing stop codons, to avoid nonsense mediated decay; having a high edit distance between each barcode; having a high edit distance between different segments of the same barcode, to allow design of multiple probes against the same barcode (with each probe being 15-50nt long); being distinct from the human and mouse transcriptome, to avoid confusion with endogenous genes. With these criteria, we generated a set of 22 barcodes and used 18 for our library, with one barcode per gene (Fig 1a schematic).

#### FISH encoding probe design

14-16 encoding probes were designed against each of the 450bp barcodes using the Stellaris Probe Designer software (LGC Biosearch Technologies). Parameters used were: Masking level = 5 (mouse), Oligo length = 19-20nt, Min. spacing length = 2nt. Each encoding probe sequence was flanked on both sides by the readout probe sequence 5’-TCTGTTTGACGCGCT-3’ with a spacer nucleotide (A) in between the readout and the encoding probe region. The concatenated sequences, and the readout probe (/5IRD800CWN/AGCGCGTCAAACAGA) were purchased from Integrated DNA Technologies (IDT).

#### 10X sequencing data processing

Sequencing results from 10X libraries were processed and demultiplexed with CellRanger-7.1.0 pipeline version (10X Genomics). An edited mouse reference genome (mm10-2020) with the 450bp barcodes added was used for alignment and generating the UMI count matrices^46^. This allows for barcode counts to be included in count matrices. Sequencing saturation of all reads and of barcode reads were calculated for all generated libraries and were >0.6 and >0.5 respectively. Filtered count matrices were analysed using the Seurat package (version 4.3.0.1) in R (version 4.3.1). Briefly, QC was performed to keep only cells with percentage mitochondrial reads below 15%, and nFeature_RNA between 200 and 7000, to exclude empty droplets, multiplets and dying cells. Counts were then normalized, scaled to 10000 transcripts per cell using the NormalizeData() and ScaleData() function respectively. Following which, Principal Component Analyasis (PCA) was run on the top 2000 most variable features, using the RunPCA() function. Clusters were identified with the FindNeighbors() function by generating a K-nearest neighbour graph with 10-16 dimensions, and clustered using the Louvain algorithm with a resolution of 0.3, using the FindClusters() function. The cells were then represented by a 2-dimensional Uniform Manifold Approximation and Projection (UMAP) graph. Lastly, coarse cell-type annotation was performed using a combination of analyzing the most highly expressed marker genes of each Seurat cluster, and expression of known canonical marker genes for each brain cell type (neurons, oligodendrocytes, astrocytes, endothelial cells, microglia). As each cluster was represented by all batches, no batch correction or integration methods were applied. For the analysis of oligodendrocyte perturbation phenotypes, oligodendrocytes expressing barcodes linked to *mSafe, Dpp6, Lrrk2, Gfap, Cfap410, Rbfox3* are grouped as “control”, as these genes are not expected to impact oligodendrocyte function.

#### Assignment of barcode/gRNA status for 10X and Stereo-seq sequencing data

For assignment of barcode-positive and barcode-negative status, negative status was assigned to cells with no expression of any barcodes, and all other cells were positive. For assignment of gRNA status, cells were classified by the expression of barcodes as follows: No barcodes expressed = Neg; more than 1 unique barcode expressed = Multiple; exactly 1 unique barcode expressed = the gene corresponding to that barcode. For oligodendrocytes, C9orf72, Clu, Lrrk2, mSafe were grouped as “control” as these genes are very lowly-expressed in oligodendrocytes and/or their KO are expected not to have an effect in oligodendrocytes. For neurons, Trem2, Olig2, Gfap, Stk39 and mSafe were grouped as “control”.

#### Gene editing analysis

Following preparation and sequencing of iSeq libraries as described above, quantification of percentage of indels were calculated with CRISPresso2 (v2.0.20) with default parameters (default_min_aln_score 60, quantification_window_center -3, exclude_bp_from_left 15, exclude_bp_from_right 15, quantification_window_size 1, plot_window_size 20, min_bp_quality_or_N 0). % Indel for each guide is calculated as % Modified (edited) - % Modified (WT). For all loci, there was at least 20k aligned reads.

#### Stereo-seq data processing

For processing of fastq files, SAW pipeline version V7.0.0 and Image Studio version 3.0.0 was used. Genome and gtf files used for alignment was generated by combining mouse reference GRCm39 and the 450bp barcodes. CID (Coordinate Identity) sequences were mapped to the coordinates on the Stereo-seq chip, allowing 1 mismatch. Reads are filtered based on Q30 phred quality score, indicating less than 1 in 1000 chance of base calling error. Following alignment to the reference, a gene count file was generated quantifying deduplicated, annotated reads. This gene count matrix is then aligned to the image (image registration). Cell segmentation was conducted using nuclear-stained tissue images and gene expression data, with the DeepCell algorithm. Following generation of a Stereopy object from the cellbin GEF file, a h5ad file and Seurat object was generated. For QC, cells with percent.mito > 20, and number of counts between <100 or > 6000 were discarded. After QC, cells were processed through the standard Seurat pipeline, as described under the “10X sequencing data and processing” methods section. Neurons expressing barcodes linked to *mSafe, Gfap, Stk39, Trem2, and Olig2* were grouped as control, as they are lowly expressed in neurons and/or are not expected to exert a strong phenotyping change.

#### Selection of hippocampal neurons with BANKSY

BANKSY was run on each chip separately, after data scaling, and before PCA using npcs = 30. We used lambda 0.2 to achieve spatially-informed cell type embeddings. Higher lambda values resulted in spatial domain segmentation with separate sections on the same Stereo-seq chip being clustered separately. The following parameters were used: spatial_mode = “knn_r”, ndim = 2, k_geom = 15. Clusters corresponding to hippocampal CA1, 3 and dentate gyrus (DG) neurons, the hippocampal neuronal niche, were kept for analysis.

#### Differential gene expression (DGE) and analysis

DGE analysis was performed with Seurat’s FindMarkers function with the following parameters were: min.pct = 0.1, logfc.threshold = 0.5, test.use = "wilcox". For reporting of the number of DEGs, the same number of cells for each perturbation was used, to ensure comparability. For calculation of neighbour DEGs, 15 neighbours were identified with BANKSY (spatial_mode = “knn_r”) for each perturbed cell in the group, and barcode-positive neighbours were excluded. The wildtype neighbours of the 2 groups (case and control), were then compared. Genes with p < 0.05 and lfc > 0.5 were counted as DEGs. For volcano plots, p values were adjusted with the p.adjust function to obtain fdr values. Genes with fdr < 0.05 and lfc > 0.5 were marked as significant.

#### Cell-cell communication (CCC) analysis

We used the LIANA (Ligand-Receptor Analysis) tool, a computational pipeline for prioritizing ligand-receptor interactions based on databases like CellPhoneDB, connectomeDB2020, and CellChat, to elucidate cell-cell communication between neurons. As cells within the same microenvironment are more likely to be interacting, we focussed on the communication between perturbed cells and their immediate neighbours. For each of these 3 perturbed groups, Srf-KO, Lrrk2-KO, and control, the perturbed cells and their wildtype/unperturbed neighbours were identified. 15 neighbours were identified for each perturbed cell with BANKSY (spatial_mode = “knn_r”), and barcode-positive neighbours were excluded. For each pair (Srf-KO + neighbours, Lrrk2-KO + neighbours, control-KO + neighbours), we applied LIANA with statistical methods “sca” and “natmi” to calculate and prioritize top interactions, as these are most likely to be biologically relevant. We modelled the perturbed cells as the source cells and their wildtype neighbours as the target cells. We then identified common ligand-receptor pairs from the prioritized list between Lrrk2-KO and control, and Srf-KO and control, and show the top 20 ligand-receptor pairs with biggest differences in sca.LRscore.

#### Calculation of R2 values

The Cell Profiler software was used to calculate the coefficient of determination (R2 value) between TdTomato protein intensity and number of FISH spots of TdTomato RNA, Barcode, Actb control, from confocal images of TdTomato protein and FISH staining. A cropped image was first analyzed using IdentifyPrimaryObjects (for analysis of TdTomato protein fluorescent intensity or FISH spots), then gridded using the DefineGrid module (200 grid squares). Next, the IdentifyObjectsInGrid module was done, followed by the RelateObjects module to relate parent objects (grid squares) to child objects (TdTomato protein fluorescent intensity or FISH spots). The pearson R2 coefficient was then computed (one point per grid area) with a fixed intercept at origin, to assess correlation between TdTomato protein fluorescent intensity and number of FISH spots.

#### Power analysis with powsimR

First, we modelled the noise in the dataset, as this affects the sensitivity of the platform. To do this, the mean-dispersion relationship was estimated using a negative binomial model, using counts data from neurons from chip2 randomly down-sampled to 10000 cells, and normalized with scran. These distribution statistics were then used to set up simulations, using the following parameters: nsims = 25, p.DE = 0.1, pLFC = 2, LibSize = “equal”. Lastly, marginal and conditional FDR and TPR are evaluated in the evaluateDE function with the following parameters: alpha.type = ’adjusted’, MTC = ’BH’, alpha.nominal = 0.1, stratify.by = ’dispersion’, filter.by = ’dispersion’, strata.filtered = 1, target.by = ’lfc’, delta = 0. As the average number of cells we obtained per perturbation was 122, marginal and conditional FDR and TPR were shown with the following number of cells: 5 vs 5, 25 vs 25, 125 vs 125, 625 vs 625.

## Data availability

Raw and processed sequencing data have been deposited at NCBI’s Gene Expression Omnibus (GEO) with accession numbers GSE274447 (Stereo-seq) and GSE274058 (scRNA-seq).

## Code availability

Analysis of the processed data in this study was done using open-source R packages Seurat (https://github.com/satijalab/seurat), BANKSY (https://github.com/prabhakarlab/Banksy), powsimR (https://github.com/bvieth/powsimR), scCustomize (https://github.com/samuel-marsh/scCustomize), and open-source Python packages Stereopy (https://stereopy.readthedocs.io/en/latest/) and SAW (https://github.com/STOmics/SAW).

## Acknowledgements

We thank Nikita Gupta, Vipul Singhal, Nigel Chou, Grace Yeo, Timothy Stuart, Adaikalavan Ramasamy and Chia Minghao for assistance and insights on bioinformatics analysis; Torsten Wustefeld for provision of Cas9 mice; Fu Yu and Christine Chiam for use of stereotaxic apparatus; Bing Shao Chia and Joshua James for assistance with library cloning; Maurice Lee and Norbert Ha for assistance with initial FISH experiments; Liew Jun Xian for cloud platform setup; Caleigh Tan for reviewing the manuscript; A*STAR’s Immunology Network (SIgN) Flow Cytometry platform for help with FACS experiments. A*STAR’s SIgN Flow Cytometry platform is supported by SIgN, A*STAR, and the National Research Foundation (NRF), Immunomonitoring Service Platform (Ref: ISP: NRF2017_SISFP09) grant. This work is supported by A*STAR Core Funding, A*STAR Central Research Fund UIBR SC18/21-1089UI and Open Fund – Young Individual Research Grant (OF-YIRG) SC18/24-727015.

## Author contributions

K.S., C.T.K. and W.L.C conceived and conceptualized the technology framework. K.S., C.T.K., W.Y.S., K.H.C. and W.L.C. designed the experiments. K.S. did stereotaxic injections, tissue collection and processing. W.Y.S. performed the FISH experiments. D.L.S, C.T.K and K.S. performed library cloning. D.L.S. and A.M. produced the AAVs. I.B.H.M. and K.S. wrote the script for generation of the barcodes. K.G. bred and maintained Cas9 mice. K.S. performed bioinformatics analysis. K.S., W.Y.S. and W.L.C. wrote the manuscript with contributions from the rest of the authors.

## Competing interests

The authors (K.S., C.T.K, I.B.H.M, W.L.C, W.Y.S) are listed as inventors on a patent application related to this work.

