## Extended Data for "Spatial Perturb-Seq: Single-Cell Functional Genomics within Intact Tissue Architecture"

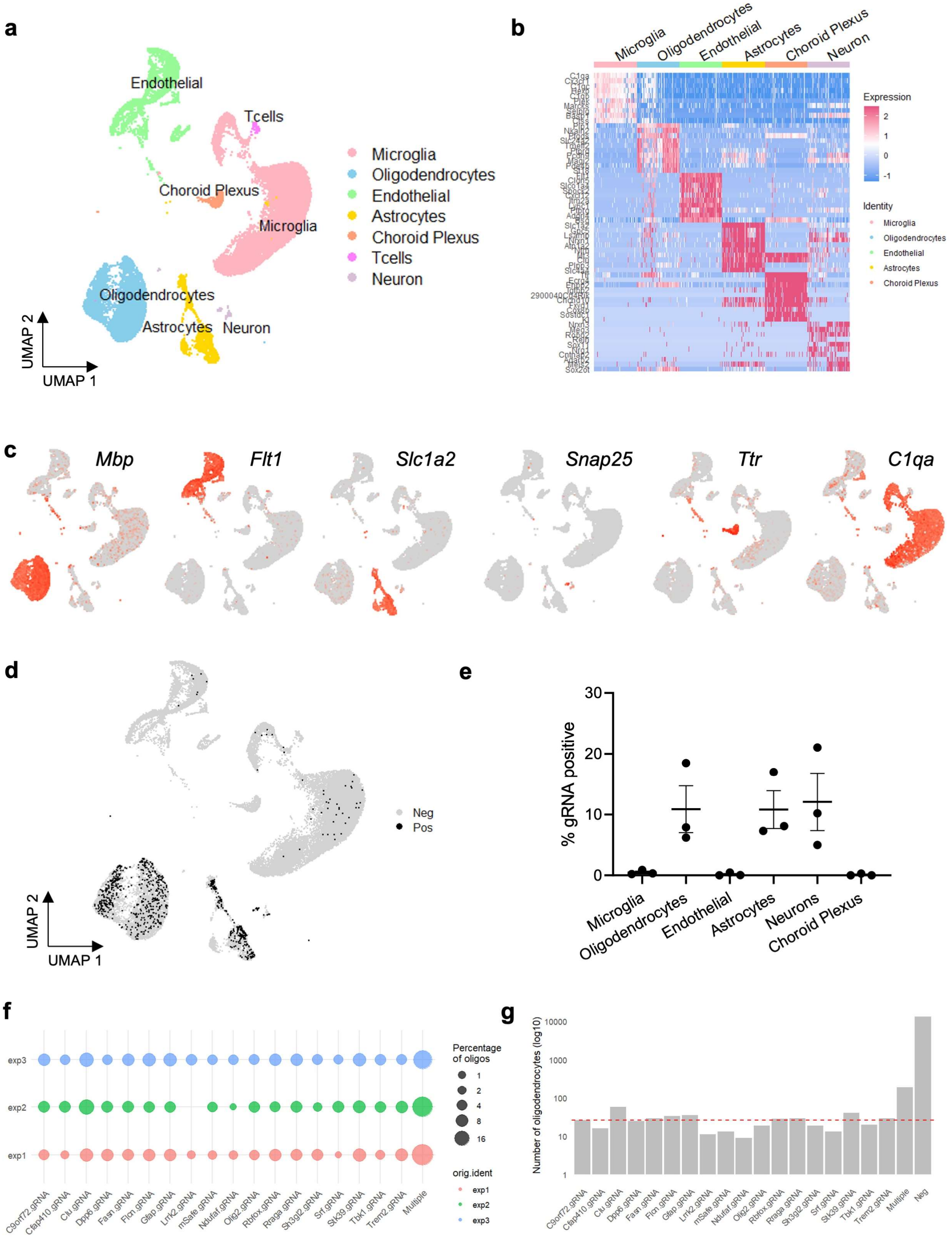


**Extended Data Fig. 1: *In vivo* perturb-seq of the adult mouse brain, with AAV-PHP.eB showing strongest tropism for neurons, astrocytes and oligodendrocytes.**

(a) UMAP projection of cells isolated from the mouse brain injected with AAV-PHP.eB, followed by 10X library prep and sequencing, showing recovery of microglia, oligodendrocytes, astrocytes, endothelial cells, T cells, neurons and choroid plexus cells. (b) Heatmap showing expression of 10 cell-type enriched genes for each cell type. (c) UMAP representation showing the expression of well-known, canonical cell-type markers for 6 major cell populations (Microglia: C1qa; Oligodendrocytes: Plp1; Endothelial cells: Flt1; Neurons: Snap25; Astrocytes: Slc1a2, Choroid Plexus: Ttr). (d) UMAP representation of all sequenced cells, showing cells with any detected gRNA barcodes in black and cells with no detected barcodes in grey. (e) Percentage of cells with detected gRNA barcodes for each annotated cell type. Each point represents a batch. Graph depicts mean and SEM. (f) Dotplot showing percentage of barcode-positive oligodendrocytes for each experimental batch. (g) Total number of oligodendrocytes for each gRNA across all 3 experiments. Min: 9, max: 58, mean: 26.

**a**

| Clu guides | Guide1 | Guide 2 | Guide3 | Guide1+2 |
|---|---|---|---|---|
| **C2C12 (cell line)** | 81% | 93% | 92% | 100% (expected 98.7%) |
| **Microglia (primary)** | 25% | 18% | 49% | 88% (expected 38.5%) |
| **Astrocytes (primary)** | 42% | 3% | 38% | 71% (expected 43.7%) |

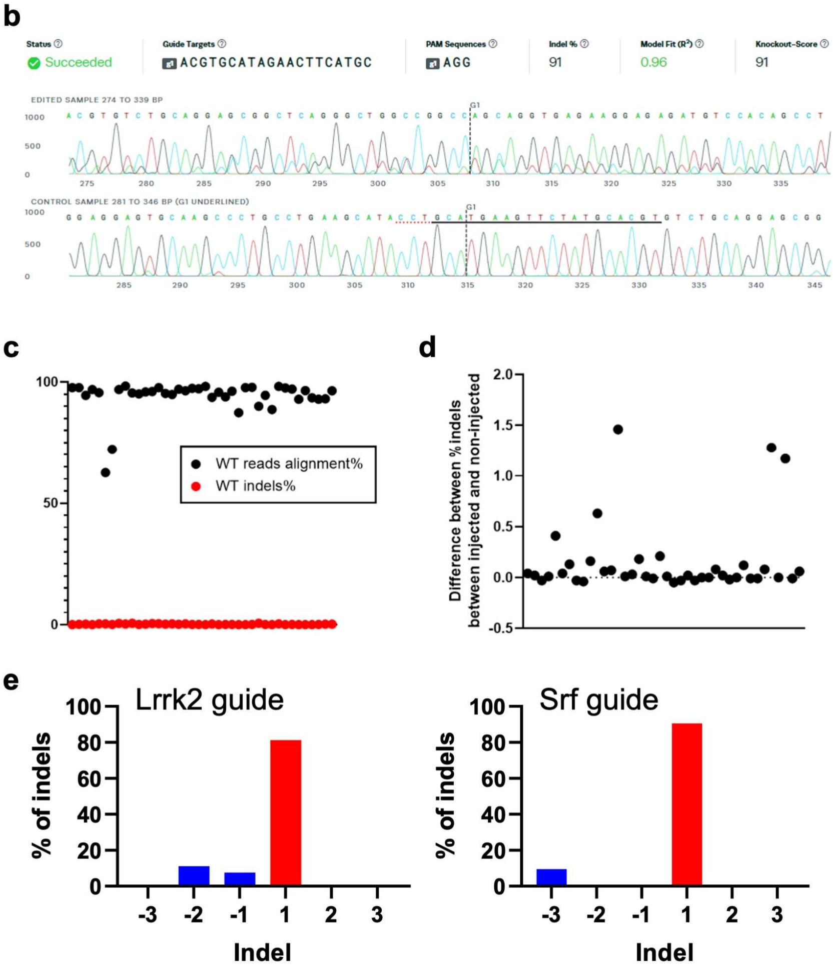


**Extended Data Fig. 2: Presence of Cas9 activity and genetic perturbations in transduced cells.**

(a) Indel% by ICE analysis, following electroporation of RNPs, using single guides and two guides in C2C12 cell line, primary microglia and primary astrocytes, showing higher than expected indel% with two guides. (b) 91% indels by ICE analysis in BMDMs from Cas9 mice electroporated with a single sgRNA, confirming Cas9 presence and activity. (c) Crispresso2 results showing good alignment of reads from control cells, and absence of indels. (d) Percentage of indels from TdTomato+ cells (with percentage of indels from WT reads subtracted). (e) Mutation profile shown for two guide RNAs, with histograms showing the distribution of indels, based on >30,000 aligned reads. No indels were detected in the corresponding un-injected control.

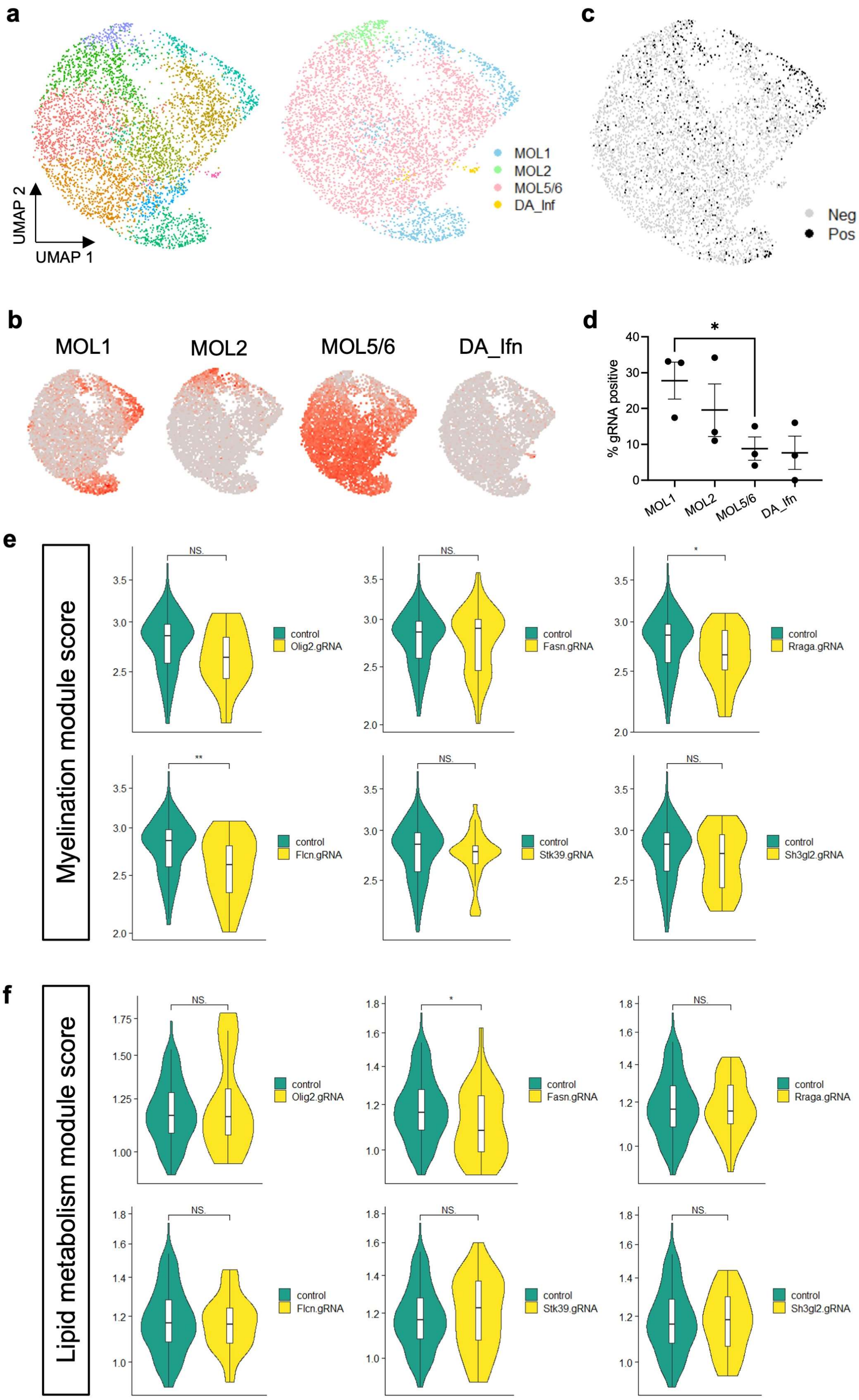
a
UMAP 2
UMAP 1
MOL1
MOL2
MOL5/6
DA_Inf
c
Neg
Pos
b
MOL1
MOL2
MOL5/6
DA_Ifn
d
% gRNA positive
*
MOL1
MOL2
MOL5/6
DA_Ifn
e
Myelination module score
control
Olig2.gRNA
Fasn.gRNA
Rraga.gRNA
Flcn.gRNA
Stk39.gRNA
Sh3gl2.gRNA
NS.
*
**
f
Lipid metabolism module score

**Extended Data Fig. 3: Analysis of perturbations in oligodendrocytes from 10X scRNA-Seq.**

(a) UMAP representation of oligodendrocytes sub-clustering, showing division into several different clusters, and recovery of different oligodendrocyte sub-types: MOL1, MOL2, MOL5/6, DA_Ifn. (b) Expression of module scores of oligodendrocytes sub-type markers (MOL1: *Fos, Egr1*; MOL5/6: *Ptgds, Opalin*; MOL2: *Klk6, Hopx*; DA_Ifn: *Ifit1, Ifit3, Stat1, Irf9*) (c) UMAP representation of oligodendrocytes, showing cells with any detected gRNA barcodes in black and cells with no detected barcodes in grey. (d) Percentage of cells with detected gRNA barcodes for each oligodendrocyte sub-type. Each point represents a batch. Graph depicts mean and SEM. (e) Violin plots showing myelination gene scores (average expression level of *Mobp, Mog, Opalin, Plp1, Mbp, Cnp, Mag, Mal*) for 6 different perturbations compared to control, showing a reduced myelination program in *Rraga*-KO and *Flcn*-KO. (f) Violin plots showing myelination gene scores (average expression level of *Srebf1, Hmgcr, Scd1, Scd2, Acaca, Dgat1, Cpt1a, Elovl6*) for 6 different perturbations compared to control, showing a reduced lipid metabolism program in *Fasn*-KO. NS, not significant; $*$ $p < 0.05$; $**$ $p < 0.01$.

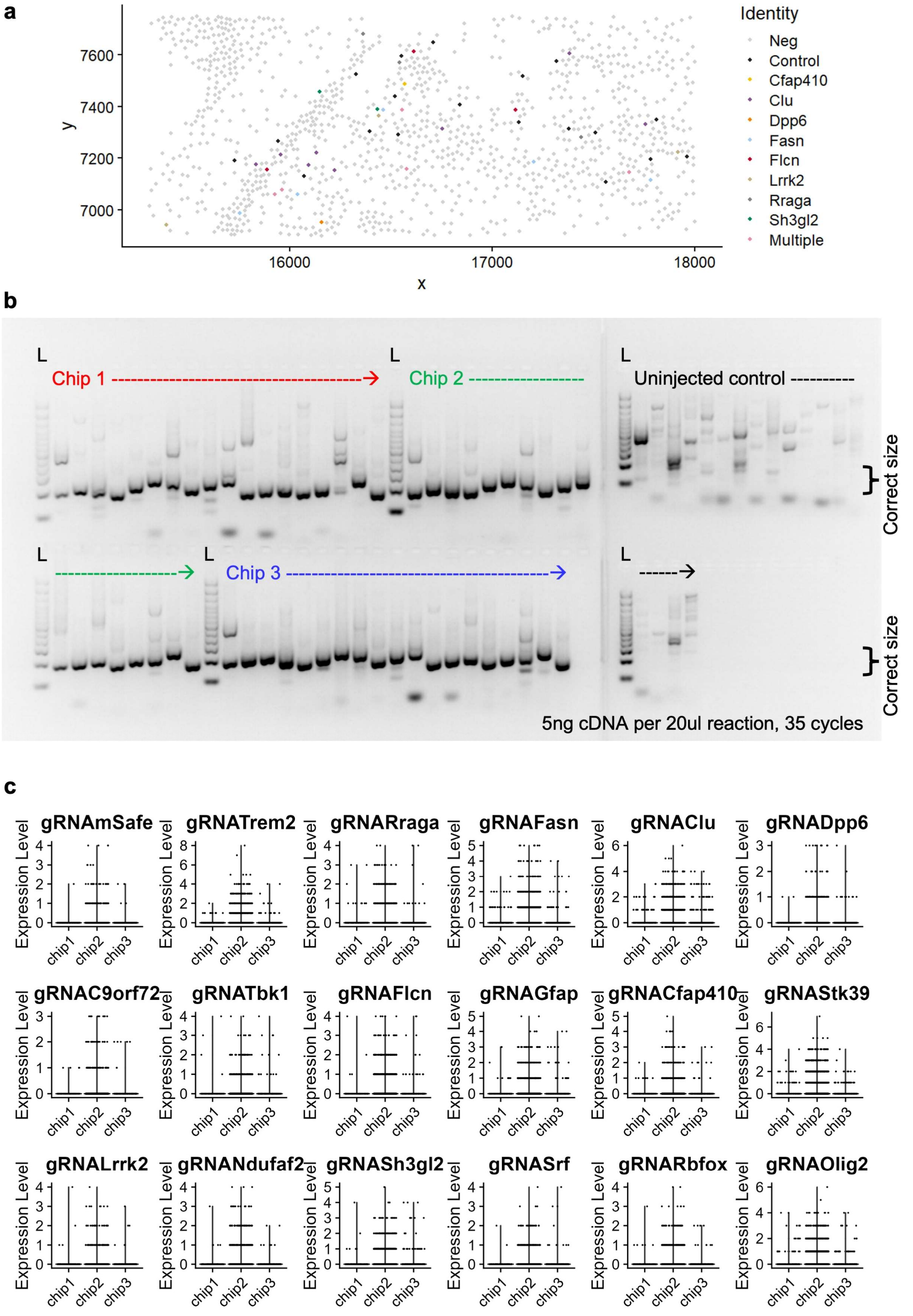


**Extended Data Fig. 4: Detection of all 18 injected exogenous barcodes in the Stereo-seq workflow.**

(a) FOV of a subsection of Stereo-seq data, highlighting individual cells, each represented by distinct barcodes shown in different colours, with cells negative for barcodes in grey. (b)

Detection of exogenous barcodes through PCR amplification of cDNA synthesized from RNA captured from chips 1, 2, and 3. All 18 barcodes were successfully detected, demonstrating effective RNA capture of exogenous sequences. (c) Expression levels of all 18 barcodes for each chip, following the whole Stereo-seq workflow.

**a**

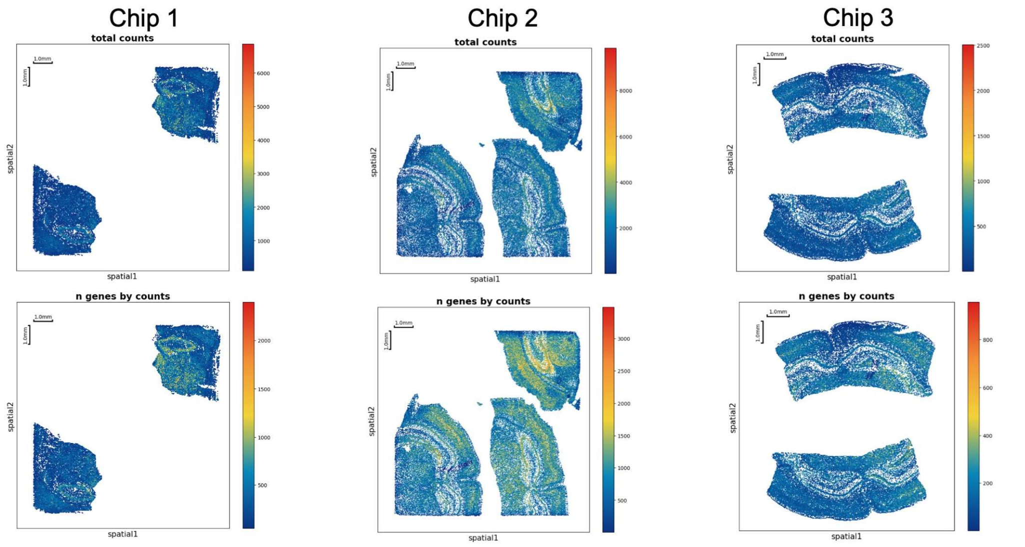


**b**

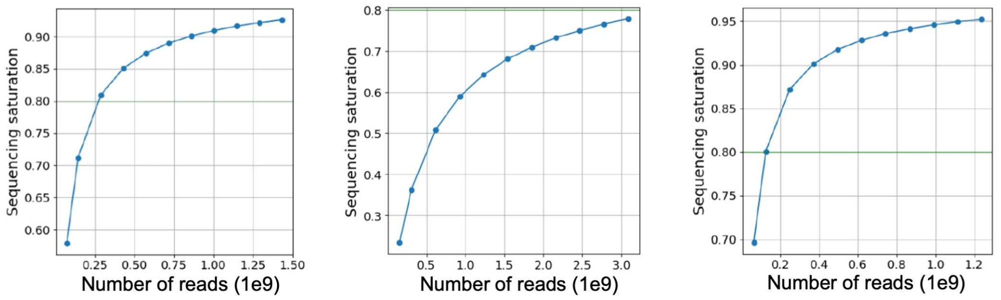


**c**

| | Total Cell counts | Mean MID counts | Mean gene counts |
|---|---|---|---|
| **Default SAW Algorithm** | 65,144 | 481.9 | 250.7 |
| **Deep Learning V1pro Model** | 63,235 | 546.4 | 281 |
| **Deepcell model** | 39,862 | 764.5 | 382.9 |

**d**

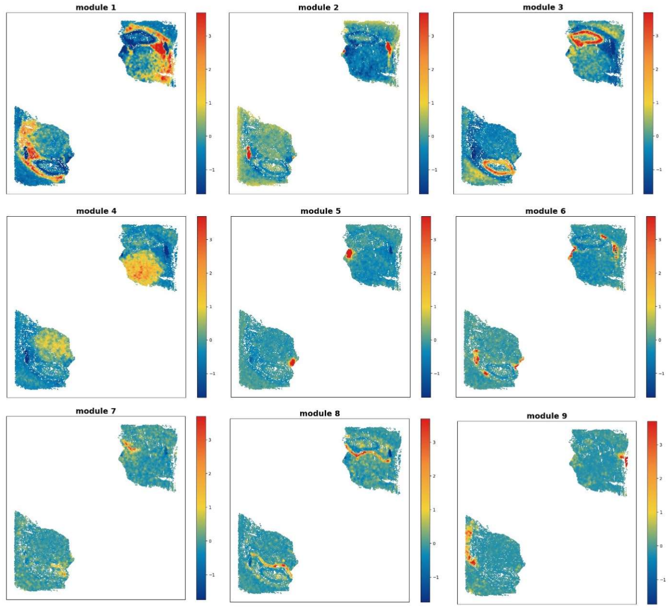


**e**

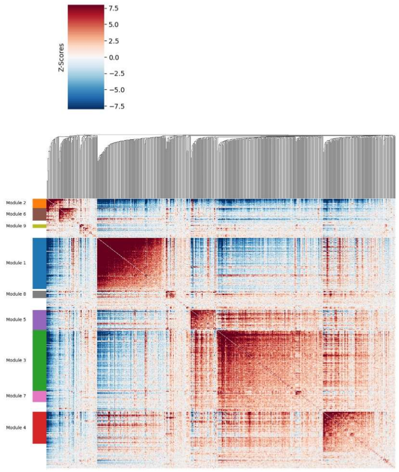

**Extended Data Fig. 5: Spatially-resolved transcriptomics of the mouse brain.**

(a) MID counts map and gene counts map following cell segmentation using the DeepCell model, for all 3 chips. Chip 1 contains 2 sections from the same mouse. Chip 2 contains 3 sections from 2 different mice. Chip 3 contains 2 sections from the same mouse. (b) Sequencing saturation curves, for all 3 chips. (c) Cell segmentation statistics using three different segmentation algorithms. Deepcell model was used for analysis. (d) Spatial hotspot analysis reveals nine gene expression modules that are consistent with both brain sections. Spatial distribution of module scores for each of the modules. (e) Heatmap depicting gene expression profiles of the nine modules. (c) – (e): Data from chip 1.

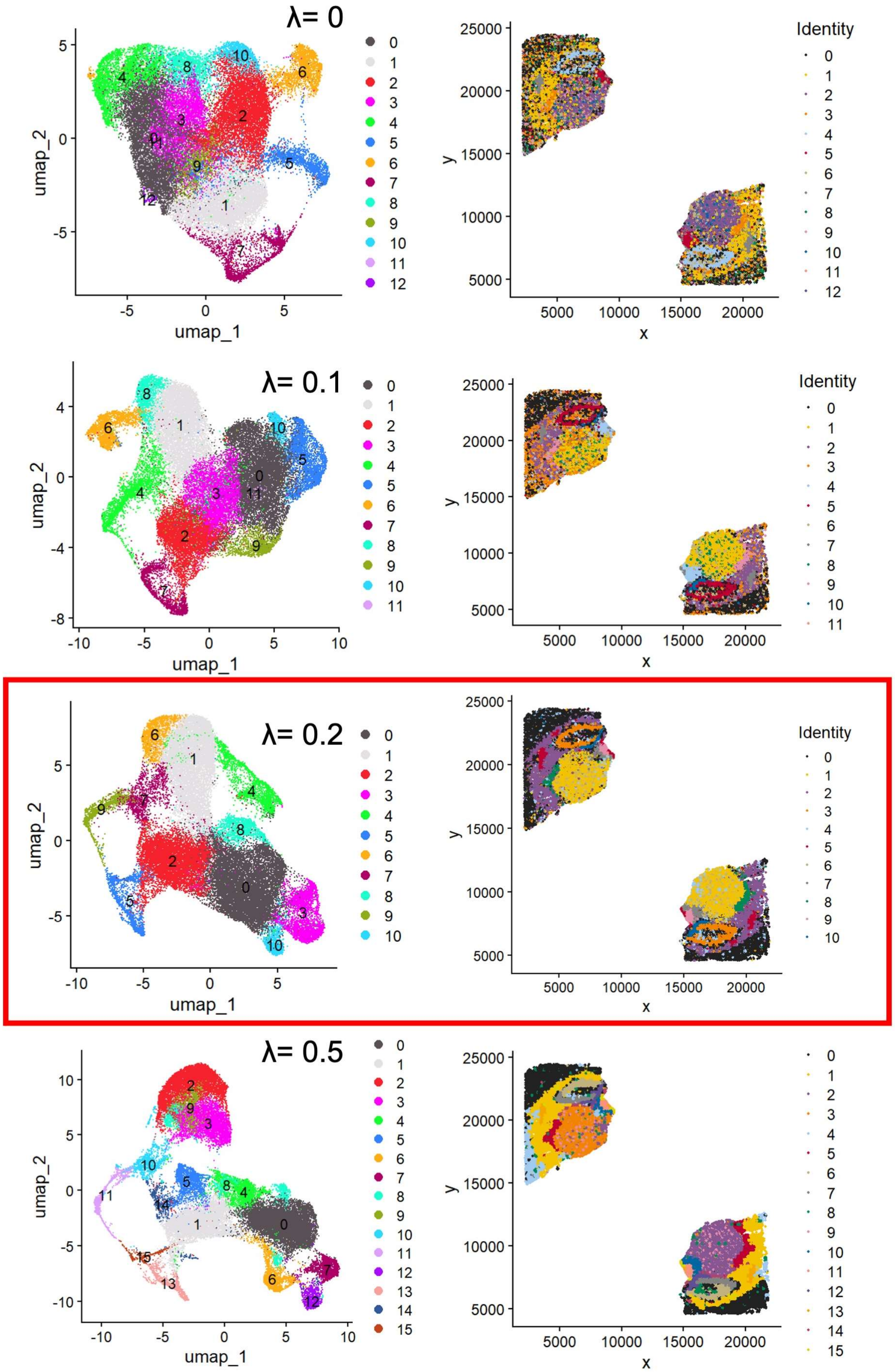


**Extended Data Fig. 6: Parameter sweep of λ used for BANKSY embedding.**

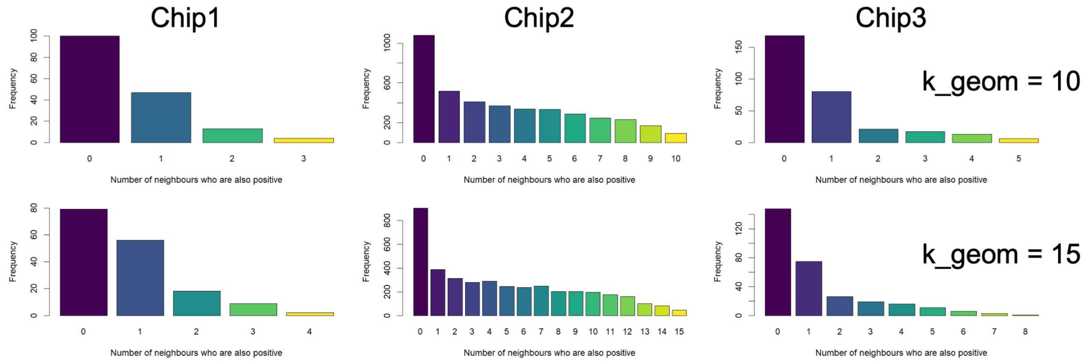


**Extended Data Fig. 7: Majority of perturbed cells have no perturbed neighbours.**

Frequency histogram of barcode-positive cells in chip 1, chip 2, and chip 3, showing the number of perturbed neighbours. Top: 10 neighbours called. Bottom: 15 neighbours called.

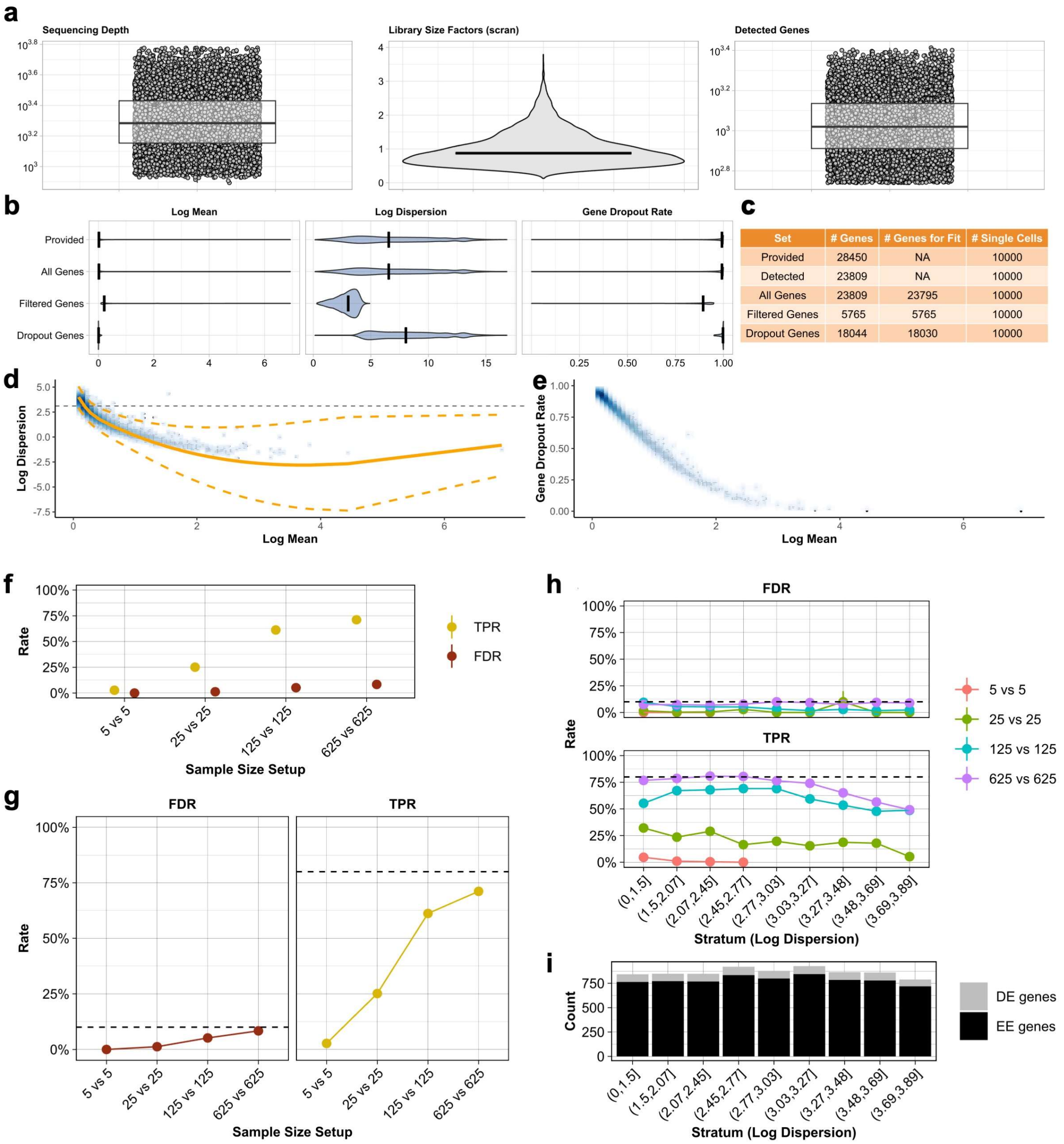


**Extended Data Fig. 8: Power analysis with powsimR.**

(a) Quality control metrics showing sequencing depth (left), library size factors (middle) and detected genes (right). Black line denotes the median. (b) Marginal Distribution of gene mean, gene dispersion and the dropout rate. (c) Number of genes and samples (cells) provided for the modeling. Detected: The number of genes and samples with one or more count. All: The number of genes for which the mean, dispersion, and dropout rate could be estimated, excluding outliers. Filtered: The number of genes above the filtering threshold, for which mean, dispersion, and dropout rate could be estimated, excluding outliers. Dropout Genes: The number of genes excluded. (d) Local polynomial regression showing the relationship between the mean and dispersion with variability indicated by the orange band. The common dispersion estimate is represented by the grey dashed line. (e) The proportion of dropouts against the estimated mean expression for each gene. (f) Marginal error rates (FDR and TPR) per sample size. (g) Marginal FDR and TPR, with dotted line indicating nominal alpha level (type I error) at 0.1, and nominal 1-beta level (type II error) at 0.8. (h) Error rates stratified by dispersion. Conditional FDR and TPR per sample size per stratum. (i) The number of equally (EE) and differentially expressed (DE) genes per stratum. For f-i, modeling is done with lfc = 2 for DE genes.

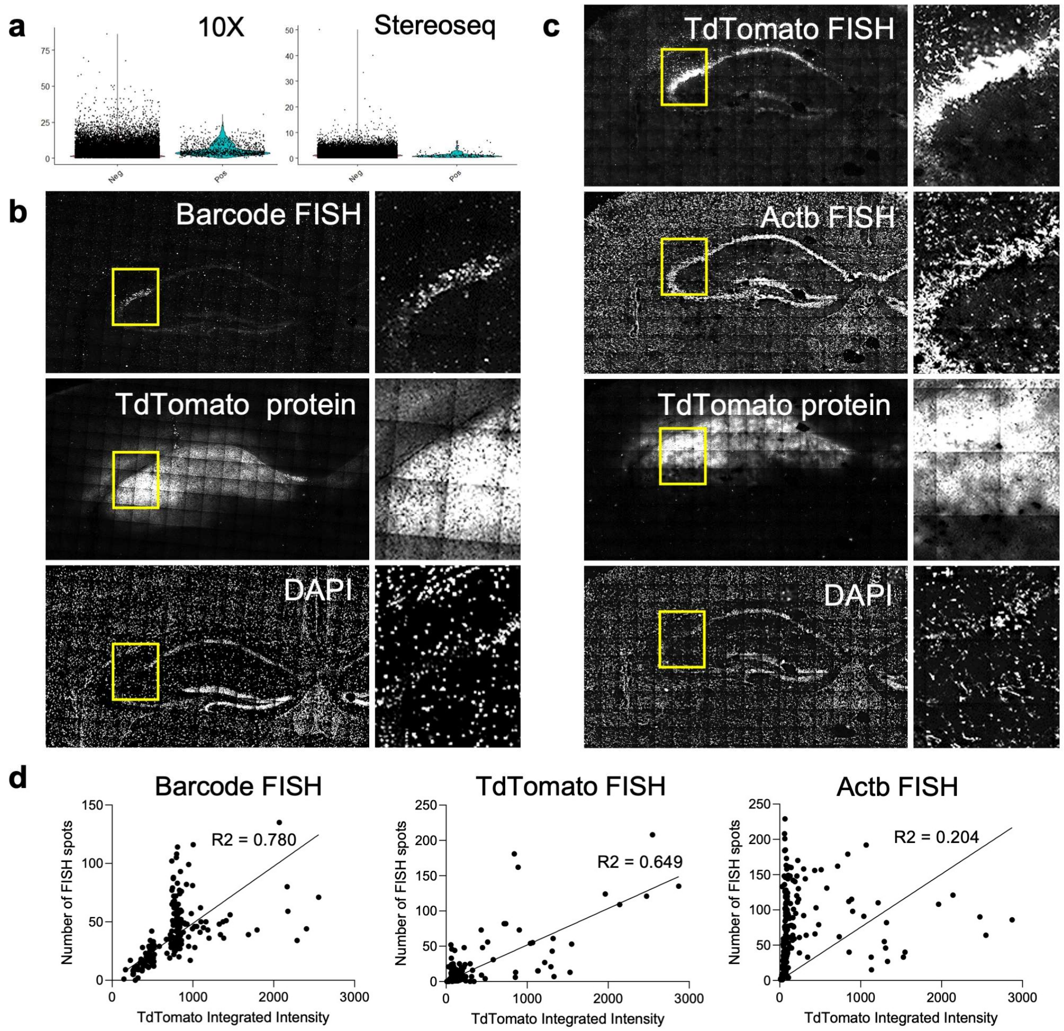


**Extended Data Fig. 9: Compatibility of Spatial Perturb-Seq with FISH.**

(a) Percentage of mitochondria reads in cells positive and negative for barcodes, showing barcodes are tolerated in cells and do not cause major deleterious effects. (b) Confocal images of the same coronal mouse brain section showing FISH staining of barcode 22 (top), TdTomato protein (middle), DAPI (bottom). (c) Confocal images of the same coronal mouse brain section showing FISH staining of TdTomato, Actb, TdTomato protein and DAPI (d) Correlation scatter plots of TdTomato fluorescent intensity against number of FISH barcode spots, FISH TdTomato spots and FISH Actb spots, showing correlation with barcode FISH and TdTomato FISH, but not Actb.
